# Cryptic genetic variation in a heat shock protein shapes the expressivity of a mutation affecting stem cell behaviour in *C. elegans*

**DOI:** 10.1101/2020.09.29.318006

**Authors:** Sneha L. Koneru, Mark Hintze, Dimitris Katsanos, Michalis Barkoulas

## Abstract

A fundamental question in medical genetics is how the genetic background modifies the phenotypic outcome of key mutations. We address this question by focusing on the epidermal seam cells, which display stem cell properties in *Caenorhabditis elegans*. We demonstrate that a null mutation in the GATA transcription factor *egl-18*, which is involved in seam cell fate maintenance, is more tolerated and thus has lower expressivity in the divergent CB4856 isolate from Hawaii than the lab reference strain N2 from Bristol. We identify multiple quantitative trait loci (QTLs) underlying the difference in mutation expressivity between the two isolates. These QTLs reveal cryptic genetic variation, which acts to reinforce seam cell fate through potentiating Wnt signalling. Within one QTL region, a single amino acid deletion in the heat shock protein HSP-110 in CB4856 lowers *egl-18* mutation expressivity. Our work underscores that natural variation in conserved heat shock proteins can shape mutation expressivity.

## Introduction

Since the completion of the human genome project, there has been a long-standing interest in using genomic data to predict key phenotypic traits. Understanding the genotype-to-phenotype relationship represents one of the goals of personalised medicine, which aims at exploiting the predictability in this relationship to determine an individual’s predisposition to disease or response to therapeutic treatment. Nevertheless, a fundamental challenge in this endeavour stems from the fact that genes exert their effects on a particular phenotype via complex gene-by-gene interactions (Chandler et al., 2013), therefore genetic variation present in the background can modify the genotype-to-phenotype mapping. Even when considering monogenic disorders, genetic modifiers can modulate the penetrance or expressivity of disease-causing loci, thereby influencing the proportion of the population developing the disease (incomplete penetrance) or the severity of the disease for each individual in the population (variable expressivity (Fournier and Schacherer, 2017)).

With regard to model organisms, mutant alleles are commonly studied in a single reference strain. Lab reference strains are highly inbred and have been selected to minimise genetic variation, which can act as confounder in the interpretation of genetic results. However, laboratory-driven evolution can fix alleles that are not widely present in the wild (Gasch et al., 2016; Sittig et al., 2016). In *Caenorhabditis elegans*, the laboratory reference strain N2 (isolated from Bristol, UK), has several adaptations suited to the lab environment (Sterken et al., 2015). For example, N2 carries laboratory-derived alleles for genes such as *npr-1*, which influences a large number of phenotypes or *nath-10*, which affects life-history traits (Duveau and Felix, 2012; Sterken et al., 2015). As a consequence, it is possible that well-studied mutations in N2 can have different phenotypic outcomes in different genetic backgrounds. Such differences can vary from subtle phenotypic effects to more substantial changes, with a key example here being genes considered essential in one *S. cerevisiae* isolate, while being dispensable for survival in another (Dowell et al., 2010). Investigating how the effects of genetic mutations may vary across genetic backgrounds is important to uncover new gene functions and understand how genetic modifiers can shape phenotypic traits.

We study here this question using seam cell development in *C. elegans* as a simplified model of stem cell patterning (Joshi et al., 2010). The seam cells are lateral epidermal stem cells that contribute to the production of the cuticle, as well as produce neuronal lineages. A number of transcription factors have been shown to play a role in seam cell development (Joshi et al., 2010). Among them, GATA-type transcription factors are evolutionary conserved regulators of proliferation and mutations in these factors have been linked to disease, such as aggressive breast, colorectal and lung cancers (Tremblay et al., 2018). In *C. elegans*, GATA transcription factors play crucial roles in the development of the gut, epidermis and vulva. One particular pair, EGL-18 and its closely related paralogue ELT-6, are thought to specify seam cell fate and are direct targets of the Wnt signalling effector POP-1/TCF (Gorrepati et al., 2013). Mutations in *egl-18* result in precocious seam cell differentiation, which in embryos has been shown to correlate with misexpression of hypodermal markers in seam cells and fusion with the hypodermis (Koh and Rothman, 2001). Furthermore, EGL-18 is required for the seam cell hyperplasia observed upon Wnt signalling hyperactivation, for example following *pop-1* RNAi which results in symmetrisation of normally asymmetric cell divisions expanding the seam cell number (Gleason and Eisenmann, 2010; Gorrepati et al., 2013).

Seam cell development is invariant in the lab reference N2 with stereotypic symmetric and asymmetric divisions during post-embryonic development giving rise to 16 cells per lateral side (Katsanos et al., 2017; Sulston and Horvitz, 1977). Over recent years, *C. elegans* has been sampled from around the globe (Cook et al., 2017), which offers exciting new opportunities to study how the genetic background affects developmental traits. We have recently shown that seam cell development is robust to standing genetic variation, with genetically divergent *C. elegans* isolates displaying a comparable seam cell number to the lab reference strain (Hintze et al., 2020). Developmental robustness can nevertheless lead to accumulation of genetic variation, which may not manifest phenotypically in wild-type, but can be revealed upon perturbation (Felix and Barkoulas, 2015; Gibson and Dworkin, 2004). This is called cryptic genetic variation and is a hidden source of variation that can influence the penetrance and expressivity of disease-causing loci, so it is linked to the emergence of complex disease (Gibson and Dworkin, 2004; Felix and Wagner, 2008; Paaby and Rockman, 2014).

Mutations in genes affecting seam cell development have been recovered and studied so far only in the N2 background. We sought therefore to investigate how the genetic background influences the phenotypic outcome of mutations disrupting seam cell development. We demonstrate here that a null mutation in the transcription factor *egl-18* results in lower expressivity, and thus higher average seam cell number, in the CB4856 strain from Hawaii compared to N2. Using quantitative genetics, we map the genetic basis of the differential expressivity of *egl-18(ga97)* mutation between N2 and CB4856. We identify a complex genetic architecture with multiple quantitative trait loci (QTLs) affecting mutation expressivity, while being cryptic for seam cell development in wild-type. We show that these QTLs act as potentiators of the effect of the Wnt signalling pathway in seam cell fate maintenance. We finally show that in CB4856 a single amino acid deletion in HSP-110, which is a heat shock 70 family member, partially explains the difference in *egl-18(ga97)* mutation expressivity between the two isolates. Our results highlight a paradigm where natural variation in a conserved heat-shock protein is a key determinant of the expressivity of a mutation influencing stem cell behaviour.

## Results

### The expressivity of a null mutation in the GATA transcription factor *egl-18* is lower in CB4856 than N2

To investigate whether the genetic background could influence the phenotypic outcome of mutations affecting seam cell development, we introduced genetic perturbations targeting known seam cell regulators in N2 into two wild isolates of *C. elegans*: the commonly used polymorphic isolate CB4856 from Hawaii, and isolate JU2007, which we had previously sampled from the Isle of Wight in the UK. Over the course of these experiments, we discovered that a null mutation in the GATA transcription factor *egl-18*, which acts downstream of the Wnt signalling pathway to maintain the seam cell fate (Gorrepati et al., 2013; Koh and Rothman, 2001), was more tolerated in the CB4856 background compared to N2 or JU2007 (Figure 1B and Figure S1A). The *egl-18(ga97)* mutation expressivity was therefore lower in CB4856, leading to a higher average seam cell count in the CB4856 population (Figure 1B). Importantly, there was no significant difference in *egl-18(ga97)* mutation expressivity between N2 and JU2007, which indicated that the underlying cause was unlikely to relate to the lab domestication of N2 (Figure S1A).

**Figure 1.**
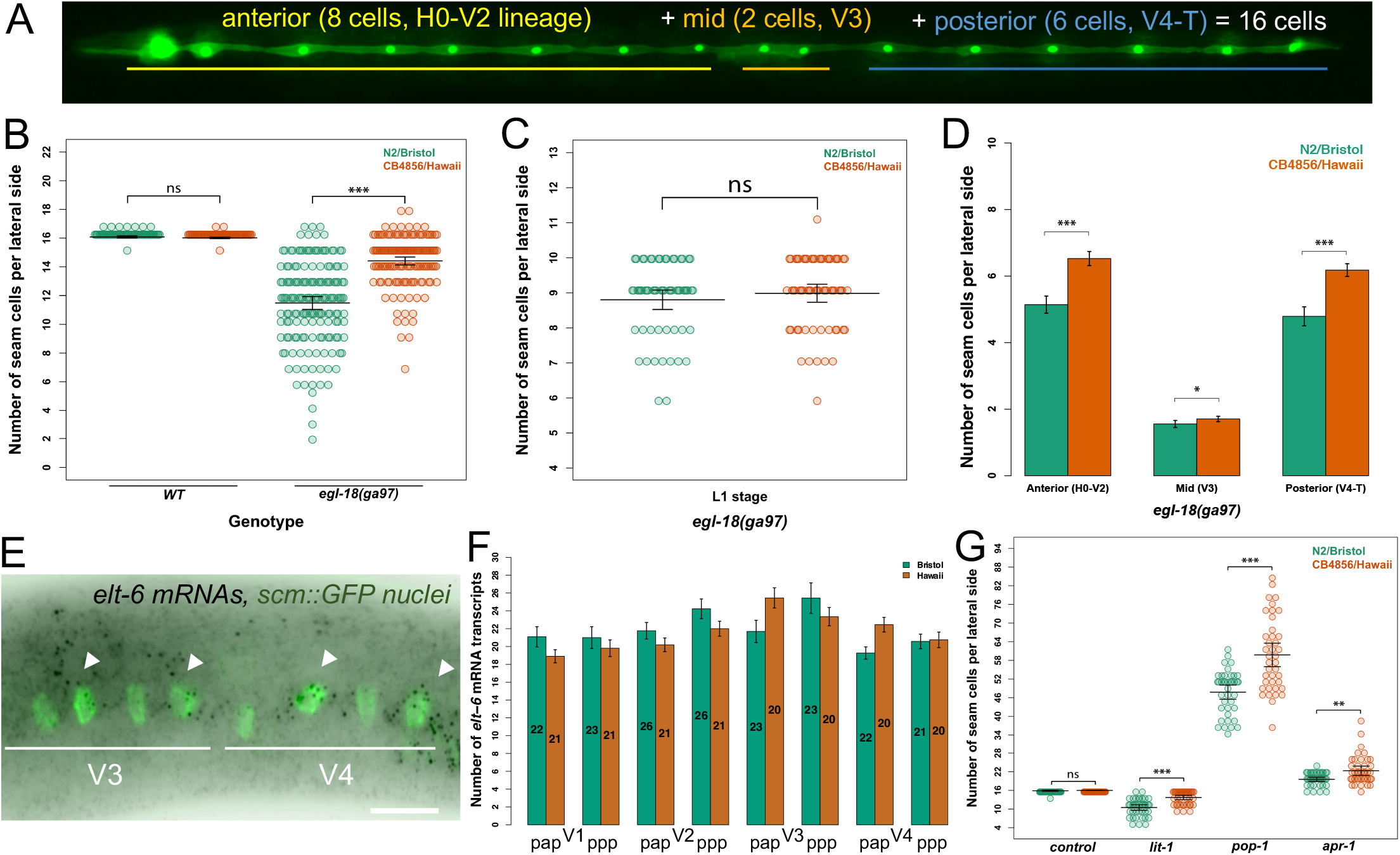
The expressivity of the *egl-18(ga97)* mutation is higher in N2 compared to CB4856 resulting in lower seam cell number in the N2 background. (A) Cartoon depicting the stereotypic seam cell distribution in wild-type animals at the end of larval development, where 8 cells are positioned anterior to the vulva (H0 – V2), 2 cells are exactly at the region of the vulva (V3), and 6 cells are found posterior to the vulva (V4 – T). (B) Average terminal seam cell number (SCN) is significantly lower in N2 than CB4856 carrying the *egl-18(ga97)* mutation (one-way ANOVA F (1, 310) = 115.02, *p* < 2.2 x 10^−16^, n ≥ 150), but is not different in wild-type (one-way ANOVA F (1, 142) = 0.1274, *p* = 0.13, n = 72). (C) No statistically significant difference was found between N2 and CB4856 after the L1 division (one-way ANOVA F (1, 122) = 0.95, *p* = 0.33, n ≥ 60). (D) Comparison of seam cell scorings of *egl-18(ga97)* mutants between N2 and CB4856 when seam cells are classified based on their position (anterior, mid, posterior). Differences in SCN in all three categories are significant between the two isolates (F (1, 310) = 66.41, 4.87, and 61.78, * *p* < 0.03 and *** *p* < 0.0001, n ≥ 150). (E) Representative smFISH image showing *elt-6* expression in posterior seam cells in an *egl-18* mutant at the late L2 stage. Seam cell nuclei are labelled in green due to *scm::GFP* expression and black spots correspond to *elt-6* mRNAs. Scale bar is 20 μm. (F) Quantification of *elt-6* expression by smFISH in N2 and CB4856 strains carrying the *egl-18(ga97)* mutation. No statistically significant difference in the mRNA was found with one-way ANOVA (F (1, 348) = 0.13, *p* = 0.72). Values within bars indicate number of cells analysed. (G) Seam cell number in N2 and CB4856 strains upon knockdown of *lit-1, pop-1* and *apr-1*. There is a significant effect of strain on SCN upon knock-down of *lit-1, pop-1* and *apr-1* RNAi, but not in empty-vector control, using one-way ANOVA (F (1, 78) = 32.71, 30.27, 7.3 and 1.37, *** *p* < 0.0001 or ** *p* < 0.001, n = 40). Error bars indicate 95 % confidence intervals.

The difference in mutation expressivity was more apparent upon strong loss of *egl-18* function, as it was less pronounced upon introgression of the milder *egl-18(ok290)* allele (Figure S1A). Perturbing other components of the seam cell network was not successful in revealing reproducible differences in seam cell number between isolates. For example, null mutations in *bro-1*, the CBFβ homologue and binding partner of RNT-1/Runx, or in the fusogen *eff-1*, led to comparable changes in seam cell number between isolates (Figure S1B-C). Therefore, we decided to investigate in more detail the genetic basis underlying the difference in *egl-18(ga97)* mutation expressivity between N2 and CB4856, two isolates for which genomic differences are well-characterised (Thompson et al., 2015; Wicks et al., 2001).

Since EGL-18 acts to maintain seam cell fate throughout development (Gorrepati et al., 2013; Koh and Rothman, 2001), the difference in the *egl-18(ga97)* mutation expressivity between N2 and CB4856 could arise from changes in embryonic or post-embryonic development. To distinguish between these two possibilities, we compared seam cell number between *egl-18(ga97)* mutants in the two isolates at the end of the L1 stage. We found that both isolates showed a comparable loss of seam cells at this early stage (Figure 1C). We then compared the spatial distribution of seam cells at the L4 stage between the two isolates carrying the *egl-18(ga97)* mutation. Here we found that *egl-18(ga97)* mutants showed a reduction of seam cells in both backgrounds, however, significantly higher seam cell counts were retained in the CB4856 background throughout the body axis (Figure 1D). There are multiple ways via which an increase in terminal seam cell number can be achieved developmentally in the *egl-18* mutant in CB4856. The first possibility involves the occurrence of additional symmetric divisions, either due to symmetrisation of normally asymmetric cell divisions, where anterior cell daughters maintain the seam cell fate instead of differentiating into hypodermis, or because of ectopic symmetric cell divisions that expand the seam cell pool. Alternatively, an increased maintenance of seam cell fate would reduce precocious cell differentiation, and thus increase terminal seam cell number independently of cell division. While phenotyping large numbers of *egl-18(ga97)* animals, we never observed tightly clustered seam cell nuclei, which are indicative of seam cell symmetrisation events (Hintze et al., 2020). These results suggest that *egl-18(ga97)* mutants show higher terminal seam cell number in the CB4856 background compared to N2 due to higher cell fate retention across all cell lineages.

One mechanism to explain mutation buffering is through redundancy within the gene network, for example if loss of function in one gene can be compensated by an increase in expression of its closely-related paralogue (Burga et al., 2011; Rossi et al., 2015; Serobyan et al., 2020). In this case, *elt-6*, could compensate for the loss of it paralague *egl-18*, and changes in the degree of compensation between CB4856 and N2 could underlie the observed difference in *egl-18(ga97)* mutation expressivity. It is of note that such genetic compensation here could only be in the form of gene expression regulation *in trans* to the *elt-6* locus, as opposed to regulation *in cis* including changes in ELT-6 protein activity. This is because *egl-18* and *elt-6* are found next to each other so the full *elt-6* locus was transferred from N2 to CB4856 together with the *egl-18(ga97)* mutation. We used single molecule fluorescent *in situ* hybridisation (smFISH) to compare *elt-6* expression between *egl-18(ga97)* mutants in N2 and CB4856 and found no statistically significant difference (Figure 1E and F). This finding suggests that the differential expressivity of the *egl-18(ga97)* mutation between N2 and CB4856 is independent of changes in *elt-6* expression.

We then hypothesised that the difference in *egl-18(ga97)* mutation expressivity between the two isolates might reflect changes in Wnt pathway activity. To test this hypothesis, we targeted genes involved in this signalling pathway by RNAi and compared seam cell counts between N2 and CB4856. We found that RNAi against lit-1/NLK decreased seam cell number in both isolates, but N2 was more sensitive to lose seam cells than CB4856 (Figure 1G). CB4856 is known to harbour variation that makes this strain insensitive to germline RNAi (Paaby et al., 2015; Pollard and Rockman, 2013; Tijsterman et al., 2002), and seam cell RNAi is also less effective in CB4856 than N2 (Figure S1D). Therefore, we cannot formally rule out that the observed difference upon *lit-1* RNAi is independent to inherent changes in RNAi sensitivity between the two isolates. Interestingly, RNAi knockdown of *apr-1/APC* and *pop-1/TCF* increased seam cell number in both isolates compared to control RNAi as expected, but this time CB4856 was more sensitive to gain seam cells than N2 (Figure S1D). This finding suggested an inherent difference between the two isolates in the outcome of Wnt-related perturbation. We compared Wnt pathway activity between wild-type N2 and CB4856 animals using an established marker reflecting POP-1 binding in cells where Wnt signalling is activated (Bhambhani et al., 2014), but found no reproducible difference (Figure S1E). Finally, the difference in mutation expressivity between the two isolates was masked upon RNAi of *elt-6* or *pop-1* (Figure S1F), highlighting a requirement for Wnt-signalling for the difference to manifest. Taken together, we conclude that the difference in *egl-18(ga97)* mutation expressivity between N2 and CB4856 is likely to reflect changes that potentiate seam cell fate maintenance by Wnt-signalled cells.

Animals carrying *egl-18(ga97)* mutations in both isolates display aberrant vulval phenotypes, such as a protruding vulva and egg-laying defects. This is not surprising because EGL-18 is required for maintenance of vulval precursor cell fate by inhibiting their fusion to hypodermis (Eisenmann and Kim, 2000; Koh et al., 2002). We tested whether the *egl-18(ga97)* mutation shows different expressivity between N2 and CB4856 using vulval cell fate induction as a read-out. We found no statistically significant difference between the two isolates (Figure S1G). We finally tested whether a previously reported polymorphism in the essential RNA cytidine N-acetyltransferase *nath-10* between N2 and other isolates including CB4856, which has been shown to influence the differential expressivity of EGF/Ras mutations in vulva development (Duveau and Felix, 2012), is able to modify seam cell number. Here, we found that the CB4856 *nath-10* polymorphism is not sufficient to modify the *egl-18(ga97)* mutation expressivity in the N2 background (Figure S1H). Taken together, these results suggested that the differential expressivity of *egl-18(ga97)* mutations between N2 and CB4856 is specific to the seam and does not rely on previously identified genetic variation that is known to affect epidermal development.

### Multiple QTLs influence *egl-18(ga97)* mutation expressivity between the two isolates

To discover the genetic basis underlying the differential expressivity of *egl-18(ga97)* mutation between N2 and CB4856, we used a quantitative genetic approach. We produced 116 homozygous recombinant inbred lines (RILs), which consisted of shuffled parental genomes and carried the *egl-18(ga97)* mutation in the background, as well as the *scm::GFP* marker to be able to visualise the seam cells (Figure 2A). Upon phenotypic characterisation of the resulting RILs, we found that the seam cell number distribution was continuous, with most RILs displaying an average seam cell number that was intermediate between the two parents (Figure 2B). We also observed transgressive segregation, with some RILs showing a seam cell number that is higher than CB4856 or lower than N2, indicating that multiple genetic loci are likely to affect this trait.

**Figure 2:**
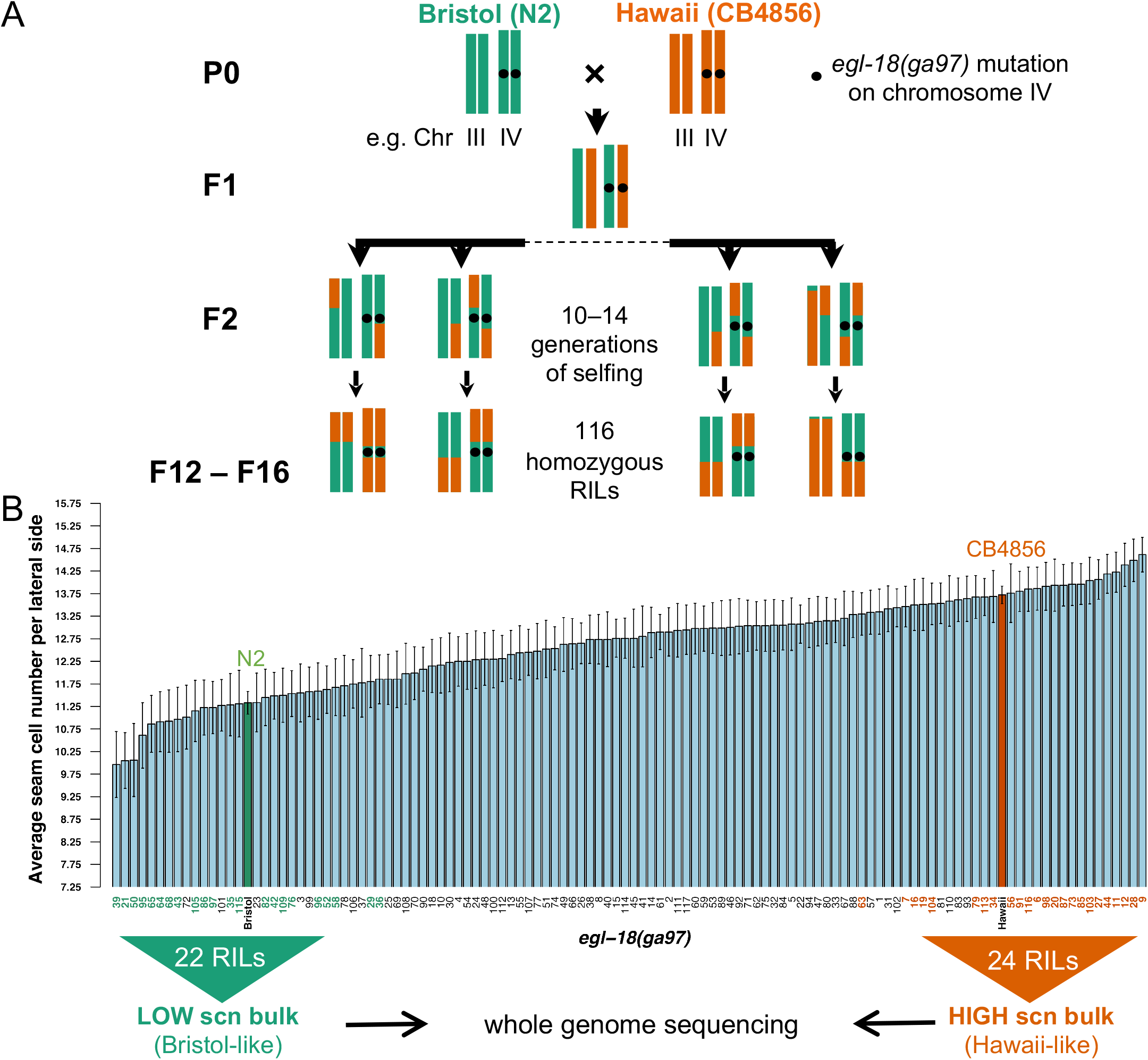
Generation and phenotypic analysis of recombinant inbred lines. (A) Generation of RILs between CB4856 and N2 carrying the *egl-18(ga97)* mutation. Two chromosomes (III and IV) are shown for simplification. (B) Seam cell number quantification in 116 RILs, ranked from lowest to highest average seam cell number. Parental strains are shown in green (N2) and orange (CB4856). The two extremes (low-bulk and high-bulk) of the phenotypic distribution were pooled for whole genome sequencing. Error bars indicate average SCN ± 95 % confidence intervals, n ≥ 77 animals per RIL.

We used bulked segregant analysis combined with whole genome sequencing to map the genetic loci involved (Frezal et al., 2018; Michelmore et al., 1991). In this approach, the extremes on each side of the phenotypic distribution that resemble the parental phenotypes (i.e. they show low average seam cell number like N2 or high average seam cell number like CB4856) are pooled and sequenced together as a bulk (Figure 2B). According to the null hypothesis, there should be no statistically significant differences in the SNP frequencies between the low-bulk and high-bulk samples for genomic regions that do not influence the expression of this phenotypic trait. However, deviations are expected for genomic regions that influence positively or negatively the phenotype. Interestingly, we observed significant deviation in SNP frequencies on multiple chromosomes, suggesting the presence of multiple QTLs (Figure 3). In particular, we identified at least four significant QTLs on chromosomes II, III, V and X that are likely to modulate the *egl-18(ga97)* seam cell phenotype (Figure 3A-F). In most cases, the high bulk contained CB4856 alleles in these QTL regions, except for the right arm of chromosome V where the high-bulk contained N2 alleles. Due to the complexity of the experimental design, impaired resolution was anticipated in certain chromosomal areas that were derived from N2 in all RILs. For example, RILs in both groups had a short region on the left arm on chromosome IV from N2, which corresponds to the region where the introgressed *egl-18(ga97)* mutation resides (Figure 3D). Also RILs in both groups had a large portion of chromosome V from N2, which is where the *scm::GFP* transgene is located (Figure 3E). Finally, a region on the left arm of chromosome I was mostly derived from N2 (Figure 3A), because this region harbours a previously known genetic incompatibility between the two parental isolates (Seidel et al., 2011; Seidel et al., 2008).

**Figure 3:**
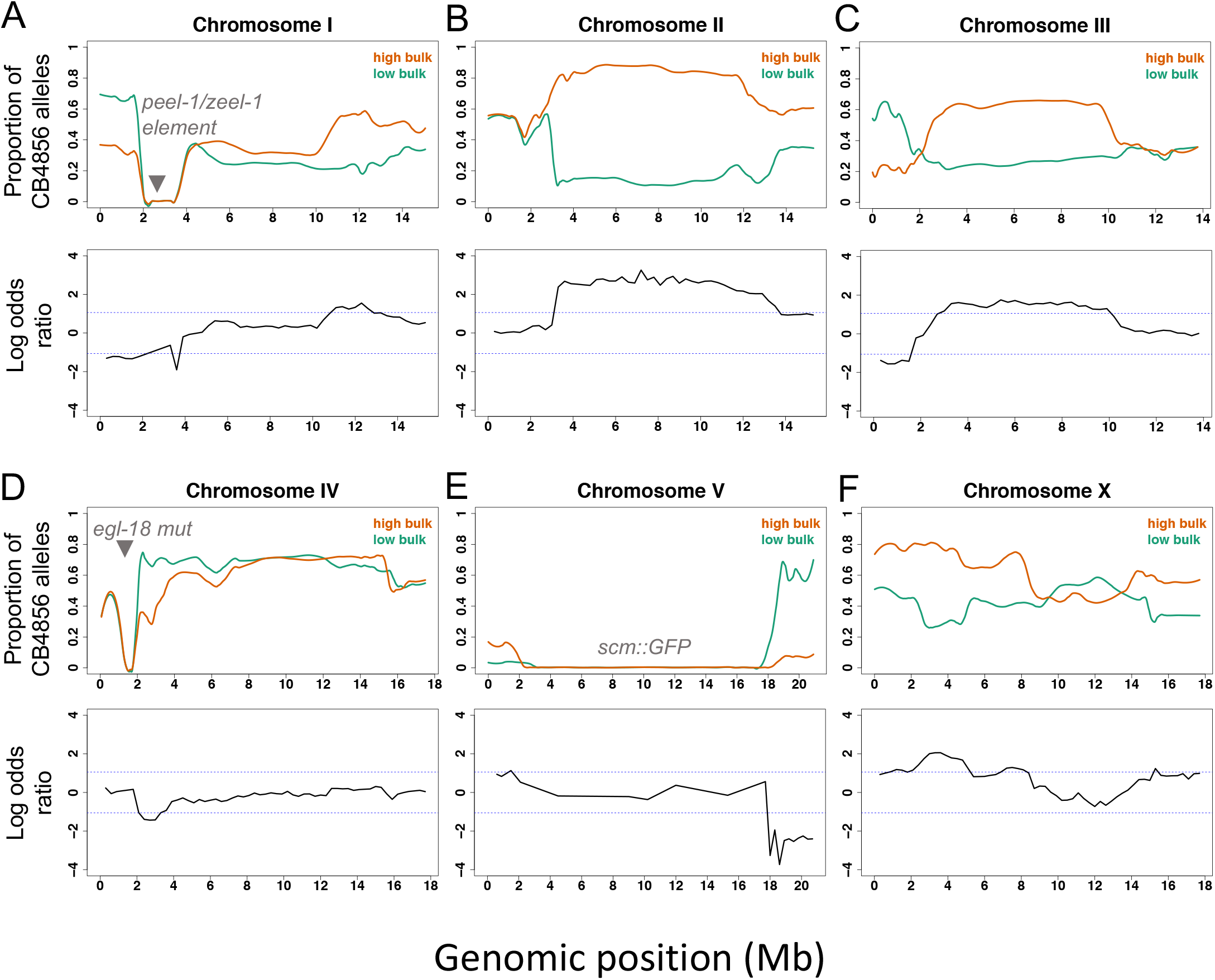
Bulk segregant analysis of recombinant inbred lines. (A-F) Proportion of CB4856 SNPs in the sequencing reads of the low-bulk and high-bulk groups is shown along the six chromosomes, from I (A) to X (F). SNP frequencies in low-bulk and high-bulk are shown in green and orange fitted curves, respectively. The curves represent locally weighted scatterplot smoothing (LOESS) regression lines from the allele frequencies at known SNP positions along the chromosomes with a span parameter of 0.1. Log-odds ratio of average SNP frequencies in non-overlapping 300 kb windows along the six chromosomes is also shown. The blue dashed lines indicate the thresholds for statistical significance for log-odd ratios at α = 0.05. The positions of the *peel-1/zeel-1* element, *egl-18(ga97)* mutation and *scm::GFP* transgene integrations are shown in grey.

To break-down the genetic composition of individual RILs in the two bulks and study their association with the seam cell number phenotype, we took advantage of known indels between the two parental strains to design markers for genotyping around the QTL regions (Thompson et al., 2015). Upon genotyping of the high and low-bulk RILS, we made the following observations. We found that 87.5% (21 out of 24) of high-bulk RILs compared to 10% (1 out of 10) in the low-bulk, carried a shared CB4856 fragment on chromosome II (4.02-7.18 Mb) (Figure S2). We also observed that 92% (22 out of 24) of high-bulk RILs compared to 40% (4 out of 10) in the low bulk, carried an N2 fragment on the right end of chromosome V (18.66 Mb – 20.99 Mb) (Figure S2). In the high bulk RILs, when chromosome II was derived from N2 rather than CB4856 (as in RILs 16 and 73, Figure S2), then CB4856 fragments were both present on chromosomes III (5.46 – 9.59 Mb) and X (3.00 - 5.08 Mb), indicating that these QTLs may act in an additive manner. RILs in the low bulk carrying large CB4856 fragments on chromosomes III and X, also contained a CB4856 fragment on chromosome V, which suggested that this chromosome V fragment may antagonise a positive interaction between the QTLs on chromosomes III and X (Figure S2). One RIL from the low-bulk (RIL-65) appeared to be the only exception in these genotype-to-phenotype correlations. This RIL contained a large CB4856 fragment on chromosome II and an N2 fragment on chromosome V, yet it displayed low average seam cell number, which highlights that additional genetic loci, beyond the identified QTLs, are likely to contribute to this complex phenotype.

### The identified QTLs harbour cryptic genetic variation affecting seam cell development through the Wnt pathway

To investigate the putative effect of these QTLs on seam cell development, we used RIL-28 to create near isogenic lines in wild type and *egl-18(ga97)* mutant N2 background. This RIL was chosen because it showed one of the highest average seam cell number in the high bulk and yet contained shorter fragments derived from CB4856 on both chromosomes II and III compared to other RILs (Figure S2). By isolating the independent fragments from CB4856 chromosomes II, III and X in N2, we found that fragments on chromosomes II and III individually increased seam cell number compared to the *egl-18(ga97)* N2 mutant phenotype (Figure 4A). The fragment on chromosome X was not sufficient on its own to increase the average seam cell number of the *egl-18(ga97)* mutant in N2, but it was able to act together with QTLs on chromosome II and III to increase the average seam cell number further (Figure 4A). Interestingly, seam cell number in lines containing a combination of two or three QTLs together rescued the mutant phenotype to CB4856 levels, suggesting that the identified QTLs capture a large proportion of the genetic variation influencing this phenotypic trait (Figure 4A). Remarkably, the same QTLs did not alter wild type seam cell development (Figure 4B), so they only acted in the presence of the *egl-18(ga97)* mutation. These results thus highlight the presence of complex genetic variation in *C. elegans*, which remains cryptic in wild-type condition but influences seam cell behaviour upon genetic perturbation.

**Figure 4:**
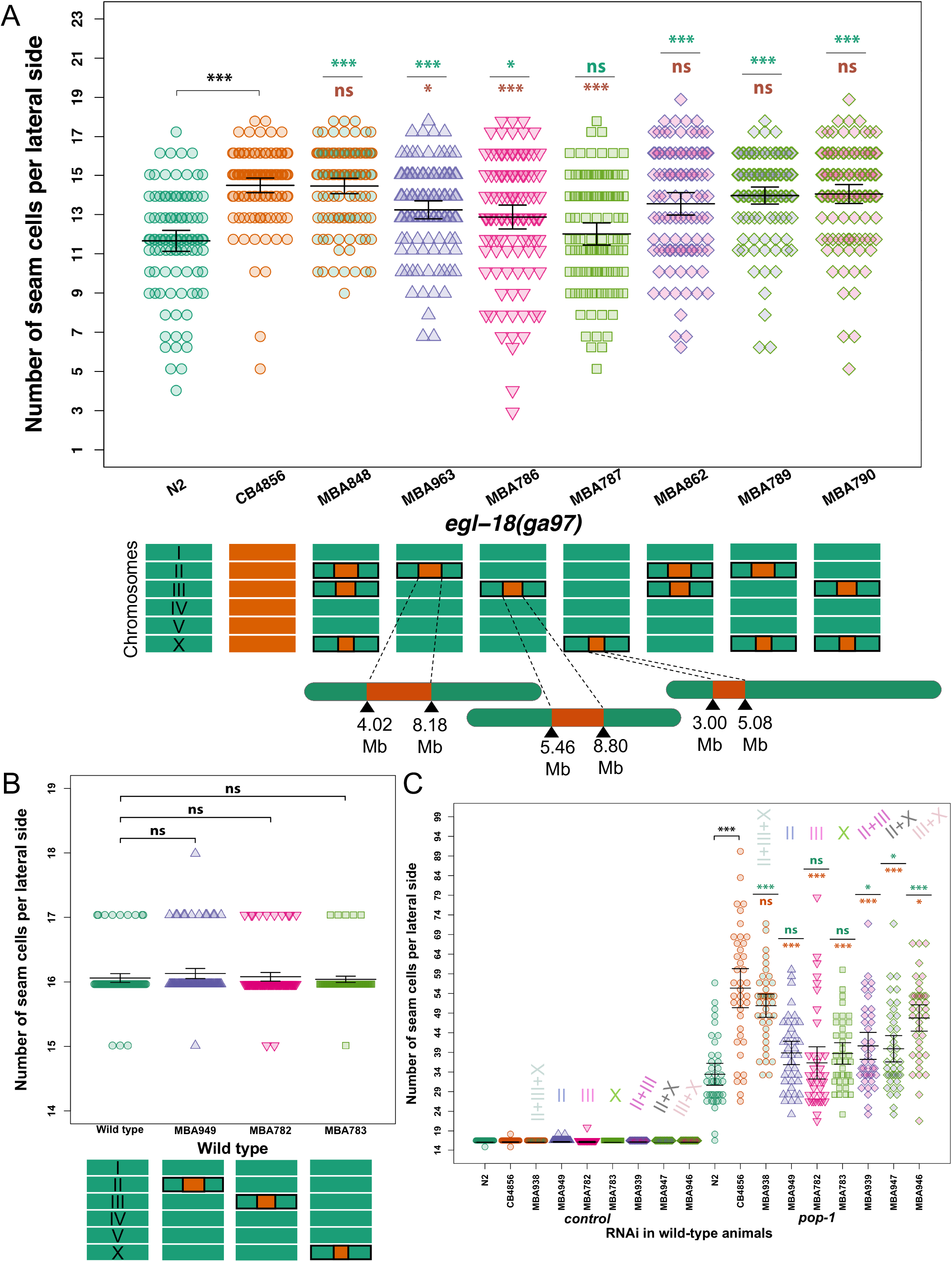
Phenotypic analysis of near isogenic lines identifies major QTLs. (A) Seam cell number in near isogenic lines containing individual and combinations of the identified QTLs from CB4856 into the *egl-18(ga97)* N2 background. One-way ANOVA showed that SCN was significantly affected by the strain (F (8, 905) = 16.55, p < 2.2 x 10^−16^, n ≥ 100). Note that QTLs on chromosomes II and III, but not on the X, were sufficient to convert the *egl-18(ga97)* mutant expressivity in the N2 background to CB4856 levels. Chromosomes below the graph depict the genotype of the strain. (B) SCN counts in near isogenic lines containing individual QTLs in a wild-type background. One-way ANOVA shows no statistically significant differences in SCN of NILs compared to wild-type with no QTLs (F (3, 396) = 1.34, *p* = 0.26, n = 100). (C) *pop-1* RNAi in near isogenic lines in a wild-type background containing individual and combinations of the identified QTLs. The same colour code is used as in panel A to define different combinations of QTLs. A significant effect of strain was found on SCN upon *pop-1* RNAi (F (8, 351) = 17.85, *p* = < 2.2 x 10^−16^, n = 40) using one-way ANOVA. Error bars indicate 95% confidence intervals and *** *p* < 0.0001 or * *p* < 0.05, correspond to significant differences by post hoc Dunnett’s multiple comparison test when compared N2 (green stars) and CB4856 (orange stars) respectively upon *pop-1* RNAi.

Given that the parental strains responded differently to knock-down of Wnt signalling components, we decided to address whether the QTLs affecting *egl-18(ga97)* mutation expressivity were related to this difference. To this end, we used RNAi to knock down the expression of *pop-1* in strains containing combinations of the QTL fragments (II, III and X) from CB4856 in a wild-type N2 background. Here we found that *pop-1* RNAi led to significantly higher seam cell number in animals carrying at least two QTL regions compared to N2. This finding highlights that the identified QTLs may promote seam cell fate by potentiating the Wnt signalling pathway.

### Natural variation in *hsp-110* contributes to the difference in mutation expressivity between the two *C. elegans* isolates

To narrow down the genomic interval of the two major QTLs on chromosomes II and III, we screened for further recombinants after crossing our NILs to N2 and screening for lines in which the presence of a defined CB4856 fragment was sufficient to increase seam cell number of *egl-18(ga97)* mutants in the N2 background. Briefly, with regard to chromosome II, we derived two NILs (strain MBA944 and MBA846), which largely rescued the expressivity of the *egl-18(ga97)* mutant in N2, while they contained mostly distinct genomic regions with a small overlapping region in the middle of approximately 0.14 Mb (Figure S3). Non-overlapping regions in MBA944 and MBA846 contained 106 and 116 genes harbouring natural variation, with 7 genes in the overlap (*dgk-5, del-10, T28D9.1, abch-1, utp-20, wrn-1, C56C10.9*). To narrow down this list further, we reasoned that genes responsible for the difference in *egl-18(ga97)* mutation expressivity between N2 and CB4856 were likely to alter seam cell number when targeted by RNAi in an *egl-18(ga97)* mutant background. However, we found no difference in seam cell number in the *egl-18(ga97)* background upon downregulating the 7 genes in the overlap (Figure S4A). These results are therefore more compatible with the existence of two independent QTLs on chromosome II, as opposed to a single QTL in the overlapping region. Consistent with this hypothesis, we found that RNAi knockdown of *egl-27* and *dsh-2*, which are located towards opposite sides of the overlap and harbour natural genetic variation known to play a role in seam cell patterning (Baldwin et al., 2016; Herman et al., 1999; Solari et al., 1999), both decreased seam cell number in the *egl-18(ga97)* mutant background (Figure S4A). Therefore, natural variation in these genes is likely to contribute to the differential mutation expressivity between the two isolates.

With regard to chromosome III, we found several loci in the broad QTL region that reduced seam cell number in the *egl-18(ga97)* mutant when targeted by RNAi (Figure S4B). We were then able to narrow down the QTL to a smaller genomic interval of approximately 1.14 Mb, which contained 59 genes in total with known genetic variation (Figure 5 and Table S3) and two strong candidates based on our RNAi experiments. First, *sor-1*, a species-specific polycomb group (PcG) protein that is involved in epigenetic silencing of Hox genes (Zhang et al., 2006). Genetic variation in this gene involved one non-synonymous substitution and one adjacent synonymous mutation, both in exon 7. The second candidate is *hsp-110*, a homolog of HSPA4 and member of the HSP70 gene superfamily (Easton et al., 2000; Nikolaidis and Nei, 2004). Genetic variation in *hsp-110* involved a 3 bp in-frame deletion in exon 5 in CB4856, which deletes a conserved amino acid within the substrate-binding domain of this chaperone (Figure 6A).

**Figure 5:**
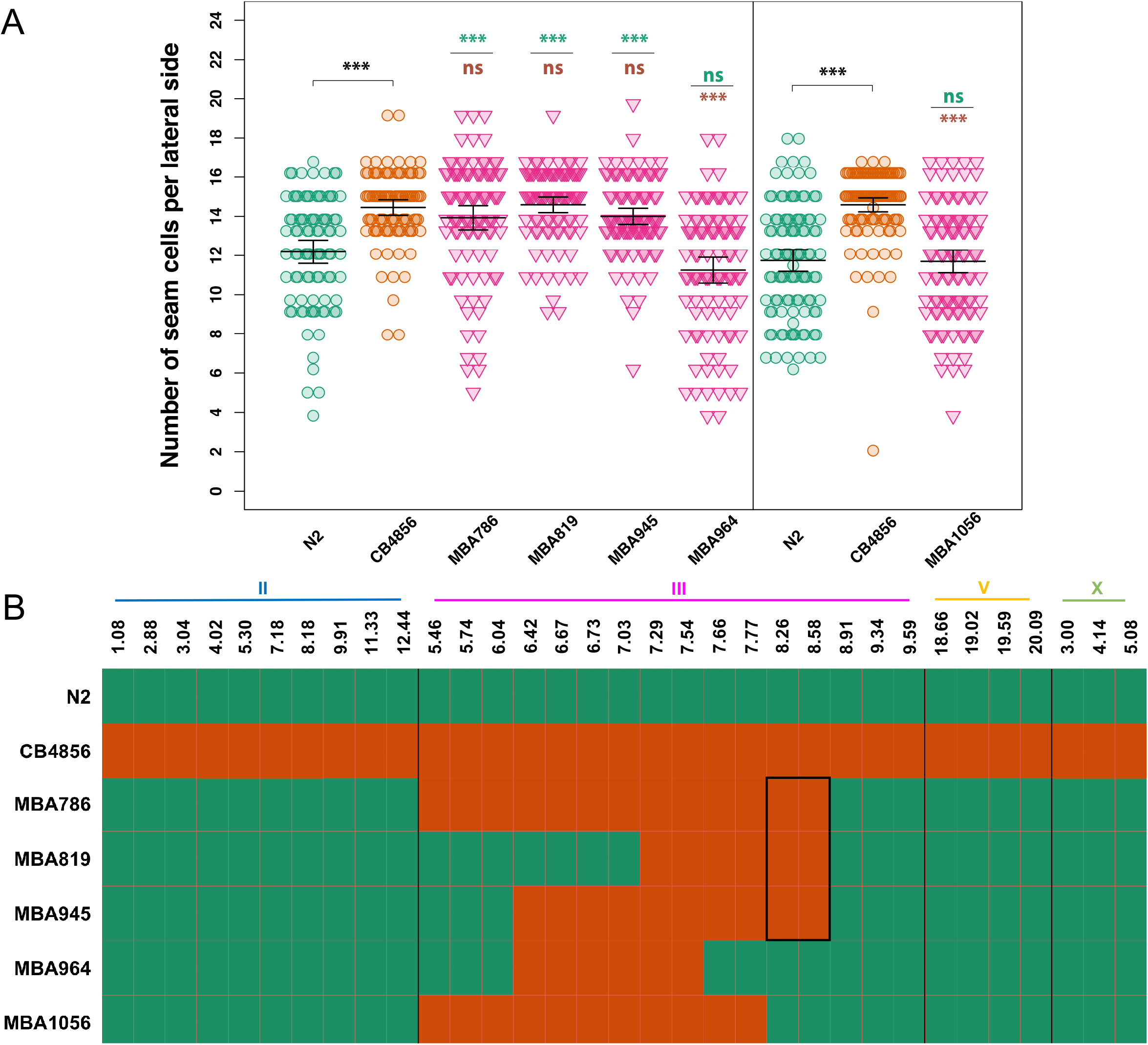
Genotype-to-phenotype analysis of NILs carrying genomic fragments of chromosome III from CB4856. (A) Seam cell number in near isogenic lines containing various fragments of chromosome III from CB4856 in the *egl-18(ga97)* N2 background. Vertical black line in the graph distinguishes between two independent experiments. In the both experiments, one-way ANOVA showed that SCN was significantly affected by the strain (F (5, 556) = 26.58, p < 2.2 x 10^−16^; F (2, 313) = 43.31, p < 2.2 x 10^−16^ respectively, n ≥ 90). Error bars indicate 95 % confidence intervals and *** p < 0.0001 corresponds to significant differences by post hoc Tukey HSD compared to N2 (green) and CB4856 (orange) respectively. (B) Genotyping of NILs carrying genomic fragments of chromosome III from CB4856 using genetic markers. Green and orange tiles represent N2 and CB4856 genomes, respectively. Highlighted black-box in the middle indicates the genomic region that is common to the NILs that convert N2-mutant expressivity to CB4856-mutant levels.

**Figure 6:**
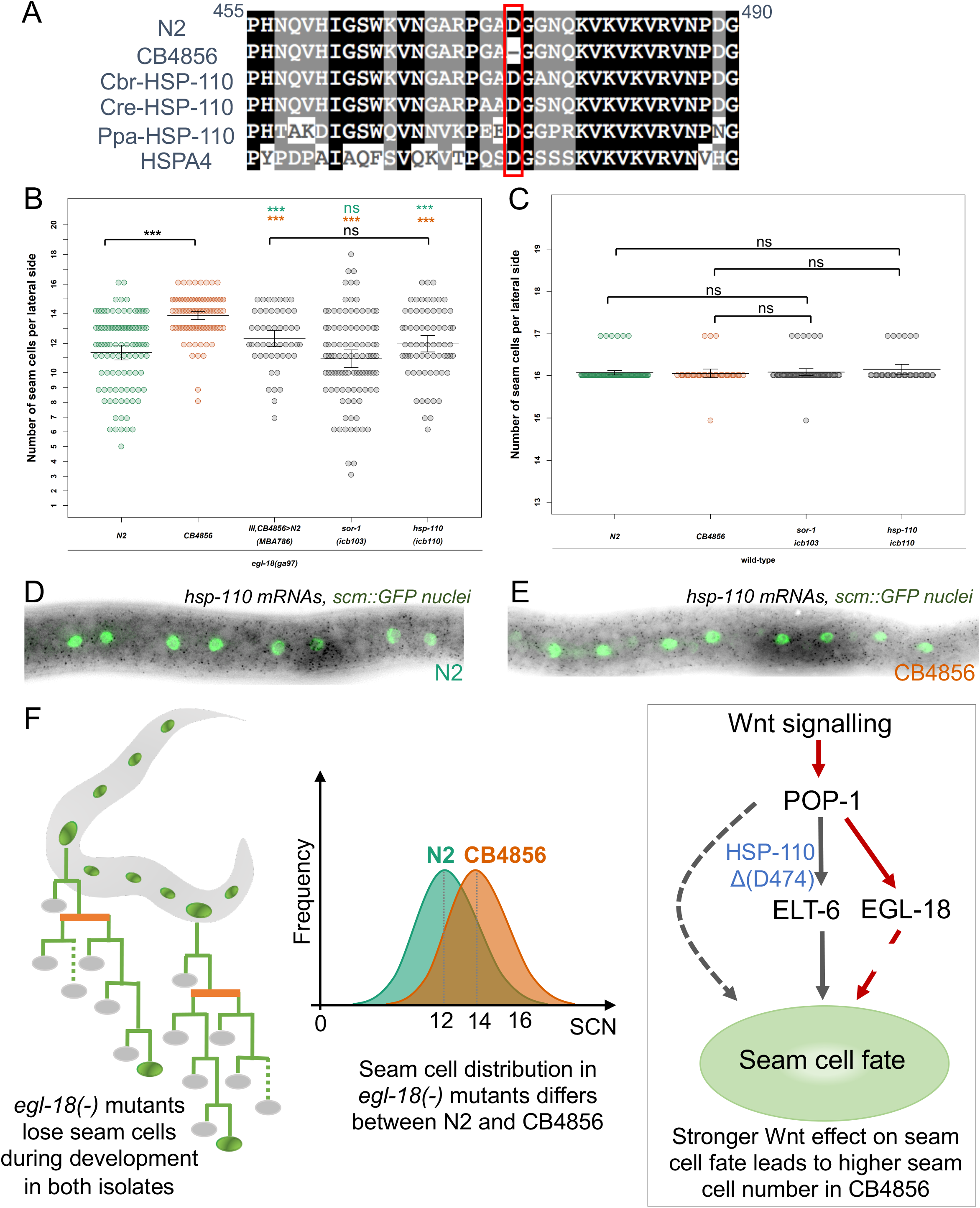
Natural variation in *hsp-110* contributes to the difference in mutation expressivity between N2 and CB4856. (A) Alignment of the HSP-110 amino acid sequence from *C. elegans, C. remanei, C. brenneri, P. pacificus* and *H. sapiens* around the amino acid deletion in CB4856 (highlighted in red). (B-C) Comparisons of seam cell counts between animals carrying the CB4856 variants in *hsp-110* and *sor-1* in an N2 background in *egl-18(ga97)* mutants (B) and wild-type (C). Comparison is shown to parental isolates with (B) or without (C) the *egl-18(ga97)* mutation, as well as the introgression of the QTL on chromosome III as a reference for the rescue. Error bars indicate 95% confidence intervals and *** *p* < 0.0001 corresponds to significant differences by one-way ANOVA followed by post hoc Tukey HSD compared to N2 (green stars) and CB4856 (orange stars) respectively. (D-E) Representative smFISH images showing *hsp-110* expression in divided seam cells in N2 or CB4856 at the late L2 stage. Seam cell nuclei are labelled in green due to *scm::GFP* expression and black spots correspond to *hsp-110* mRNAs. (F) Developmental model explaining the difference in *egl-18(ga97)* mutation expressivity between N2 and CB4856. In Wnt-signalled cells, POP-1 acts as a transcriptional activator leading to adoption of seam cell fate through the expression of transcription factors, such as EGL-18 and ELT-6. EGL-18 is a major seam cell promoting factor, so loss of seam cell fate occurs in the *egl-18(ga97)* mutant in both isolates due to differentiation of seam cells. However, in CB4856, seam cell fate is better retained in the absence of EGL-18 because of genetic variation that acts to reinforce seam cell fate acquisition via potentiating the effect of Wnt signalling through ELT-6 and/or other yet unknown factors.

To validate the impact of these variants, we used CRISPR-mediated genome editing to engineer them into N2 and assess their role on seam cell development both in a wild-type and in an *egl-18* mutant background. Interestingly, we found that CB4856 genetic variation in *hsp-110* significantly increased seam cell number when introduced into an *egl-18(ga97)* mutant background in N2 (Figure 6B), while no effect was seen in a wild-type background (Figure 6C). We used smFISH to study the expression of *hsp-110* and found evidence for expression around seam cell nuclei, with no apparent difference between N2 and CB4856 (Figure 6D-E). This indicates that HSP-110 is likely to act cell-autonomously to modify seam cell behaviour. Instead, we observed no effect on seam cell number when we introduced the CB4856 *sor-1* polymorphisms in a wild-type or *egl-18* mutant in the N2 background (Figure 6B-C). Taken together, these results suggest that natural genetic variation in a conserved heat-shock protein contributes to the difference in *egl-18(ga97)* mutation expressivity between CB4856 and N2.

## Discussion

Previous work has shown that the phenotypic effect of mutations can vary depending on the genetic background in various models (Chow et al., 2016; Dworkin et al., 2009; Sittig et al., 2016). For example, the severity of various null mutations in mice depends on the genetic background, with often opposing effects observed among strains (Sittig et al., 2016), indicating that the genetic background can influence the genotype-to-phenotype relationship in a complex way (Chandler et al., 2017). *C. elegans* has low genetic diversity due to its self-fertilization, however the genetic variation present in the background has been shown to influence the outcome of genetic perturbations, such as RNAi-driven phenotypes (Paaby et al., 2015; Vu et al., 2015), although the exact molecular changes in the genetic background that influence developmental outcomes remain poorly understood.. With regard to mutation expressivity, a lab evolved non-synonymous polymorphism in *nath-10* between N2 and the AB1 isolate from Australia has been shown to contribute to the expressivity in a mutation in the epidermal growth factor receptor gene involved in vulval development (Duveau and Felix, 2012).

We find here that the expressivity of the *egl-18(ga97)* mutation is lower in CB4856 compared to the lab reference strain N2, with approximately two more seam cells produced on average in the CB4856 background. GATA factors are important players in animal development and mutations in these factors are known to lead to blood and heart disease, with often variable phenotypic outcomes in human patients (Brambila-Tapia et al., 2017; Tremblay et al., 2018). The difference in *egl-18(ga97)* mutation expressivity is specific to the seam and manifests during post-embryonic development through changes that reinforce seam cell maintenance in a Wnt-dependent manner. Using a quantitative genetic approach, we discover at least four QTLs that modify seam cell number in an *egl-18(ga97)* mutant background, acting both independently, as well as in combination. We pin down the molecular basis of one of these QTLs and demonstrate that a deletion of a single amino acid (D474) in the heat shock protein HSP-110, which has occurred during the evolution of the CB4856 strain, is sufficient to lower mutation expressivity and thus increase the average seam cell number when introduced into N2. HSP110 proteins form a distinct branch of the Hsp70 gene superfamily (Dragovic et al., 2006; Easton et al., 2000; Raviol et al., 2006), which includes conserved proteins that assist in protein folding and protect cells from stress (Mayer and Bukau, 2005). HSP110 members are thought to assist in solubilization of protein aggregates through nucleotide exchange activity rather than acting as canonical ATPase chaperones (Rampelt et al., 2012; Scior et al., 2018). This protein has not been studied before in the context of *C. elegans* development. Previous reports have demonstrated a role for HSP-110 in lifespan upon heat shock (Rampelt et al., 2012), as well as recovery from thermal stress in plant pathogenic nematodes, where evolution in *hsp-110* has occurred in *Globodera* species through gene duplication (Jones et al., 2018).

Molecular chaperones, like Hsp90, can act as phenotypic capacitors that modulate the genotype-to-phenotype relationship (Zabinsky et al., 2019). Hsp90-mediated buffering of protein folding has been thought to influence the variable expressivity of genetic disease (Karras et al., 2017). In the seam, RNAi knockdown of *hsp-90* or DnaJ chaperones leads to increased variability in seam cell number (Hughes et al., 2019; Katsanos et al., 2017). Differences in heat shock protein expression have also been shown to affect the incomplete penetrance of mutations (Burga et al., 2011; Casanueva et al., 2012). Our work highlights that natural variation in conserved heat shock proteins can shape mutation expressivity. It is of note that mutations in HSP-110 in human patients correlate with variable responses to drug chemotherapy (Collura et al., 2014), so it is conceivable that such mutations may influence the phenotypic outcome of disease-causing loci.

EGL-18 and ELT-6 transcription factors act redundantly to promote seam cell fate (Gorrepati et al., 2013). In both CB4856 and N2, EGL-18 appears to be the major seam cell promoting factor based on the severity of the loss of function phenotype, while ELT-6 can compensate to some extent for the absence of EGL-18 activity. Genetic compensation has been shown to occur in various organisms including *C. elegans* as a mechanism to buffer against null mutations to protect from developmental failure (Burga et al., 2011; Rossi et al., 2015; Serobyan et al., 2020). However, we did not find changes in *elt-6* expression in an *egl-18* mutant background between the two isolates. We therefore propose a model wherein changes in the seam cell regulatory network through accumulation of natural variation, including the amino acid deletion found within HSP-110 in CB4856, may potentiate the Wnt signalling pathway to promote seam cell fate through ELT-6 or another factor (Figure 6F). This model of reinforced seam cell fate adoption in CB4856 through Wnt signalling is consistent with the increase in seam cell counts observed in a *pop-1* RNAi treatment in CB4856 compared to N2. Interestingly, a recent study revealed cryptic genetic variation in the requirement of the POP-1 input in gut differentiation (Torres Cleuren et al., 2019), suggesting that natural variation may more broadly shape Wnt-dependent gene regulatory networks in *C. elegans*. It is of note that a crosstalk between Hsp70 proteins and Wnt signaling has been proposed in other stem cell contexts (Zhang et al., 2016). Based on the position of the deleted amino-acid in CB4856 within the substrate binding domain of HSP-110, we speculate that some altered interaction with a protein partner may increase the strength of the Wnt signaling pathway in promoting the seam cell fate.

The amino acid deletion in *hsp-110* is specific to the CB4856 isolate of *C. elegans* and is not present in N2, JU2007 and other divergent isolates (CeNDR release 20180527). CB4856 belongs to a group of highly polymorphic strains found around geographically isolated Hawaiian islands. These isolates are thought to represent ancestral genetic diversity because they contain approximately three times more diversity than the non-Hawaiian populations (Andersen et al., 2012; Cook et al., 2017; Crombie et al., 2019). Interestingly, we previously reported a genotype-by-environment interaction affecting seam cell number (Hintze et al., 2020). An increase in temperature, from 20 to 25 °C, was reported to increase seam cell number through a symmetrisation of normally asymmetric divisions at the L4 stage in many *C. elegans* isolates, except for CB4856 (Hintze et al., 2020). It is therefore intriguing that CB4856 appears to harbour loci that both reinforce seam cell fate maintenance and suppress ectopic seam cell fate acquisition, which highlights that genetic variation in *C. elegans* can influence developmental phenotypes in a complex manner. It is of note that natural variation exists in *hsp-110* and other heat-shock proteins including *hsp-90* (CeNDR), so it remains to be seen whether such variants influence developmental outcomes more broadly.

Although natural variation in *hsp-110* contributes to the difference in mutation expressivity, we show that a complex genetic architecture underlies this phenotypic trait so other loci are also involved. Our results suggested two strong candidates for the QTL on chromosome II, that is the Dishevelled protein *dsh-2* and *egl-27*. There are three Dishevelled proteins (MIG-5, DSH-1 and DSH-2) in *C. elegans*. DSH-2 and MIG-5 have been shown to act redundantly to regulate seam cell fate by positive regulation of nuclear SYS-1 β-catenin levels (Baldwin et al., 2016). EGL-27 is an ortholog of human RERE (arginine-glutamic acid dipeptide repeats protein) and mutations in *egl-27* have been associated with seam cell defects leading to an increase in seam cell number (Herman et al., 1999; Solari et al., 1999). Chromatin-modifying regulators, such as EGL-27, have been proposed to represent highly connected hubs in *C. elegans* genetic interaction networks (Lehner et al., 2006), thereby influencing the phenotypic outcomes of multiple mutations in factors belonging to several unrelated pathways. Therefore, it is conceivable that natural variation in the core Wnt signalling component *dsh-2* and *egl-27* may be involved in the modification of the phenotypic outcome of the *egl-18(ga97)* mutation in the seam. Future work is required to understand the phenotypic consequences of these natural variants and dissect how they may modify the Wnt signalling pathway.

## Materials and methods

### Strains and genetics

*C. elegans* was maintained on a lawn of *Escherichia coli* strain OP50 seeded on Nematode Growth Medium (NGM) according to standard procedures (Stiernagle, 2006). *C. elegans* larvae were synchronised by bleaching gravid hermaphrodites and washing the eggs twice with M9 buffer or by puncturing gravid animals with an injection needle to release eggs in the case of *egl-18* mutants. All strains used in this study are listed in Table S1. Mutations in *egl-18* were introgressed into wild-type strains MBA256 and MBA19 (CB4856 and JU2007 respectively carrying the *wIs51* transgene) (Hintze et al., 2020) in a two-step cross. In the first step, MBA256 and MBA19 males were crossed to *egl-18(-)* hermaphrodites carrying the *wIs51* transgene. In the second step, F1 males from the previous cross were crossed to wild isolate hermaphrodites. F2 animals from the second cross that were egg-laying defective were isolated and the two-step cross was repeated five times to produce 10 × backcrossed strains. The process was repeated for the introgression of *eff-1(icb4)* into the MBA19 background. The *eff-1(icb4)* mutation will be described elsewhere (Koneru et al. in preparation).

### RNA interference (RNAi)

RNAi was performed by feeding animals with bacteria expressing double-stranded RNA. Bacterial clones originate from the Ahringer and Vidal libraries, or were custom made (*wrn-1, utp-20*). Bacterial clones were grown overnight in liquid LB medium with 50 μg/ml ampicillin and 12.5 μg/ml tetracycline. The bacterial cultures were seeded onto filter-sterilised isopropyl β-D-1-thiogalactopyranoside (IPTG)-containing RNAi plates and allowed to dry for 2 – 4 days at room temperature before use. Custom RNAi clones were made by amplifying gene fragments for a gene of interest using primers that contained the following sequences (Fw 5’-AGA CCG GCA GAT CTG ATA TCA TCG ATG-3’, Rev 5’- TCG ACG GTA TCG ATA AGC TTG ATA TCG - 3’) to allow Gibson cloning into EcoRI-digested L4440.

### Phenotypic analysis and microscopy

Seam cell number was quantified in day-1 adults or late L4s. Animals were anaesthetised using 100 μM sodium azide and mounted on a 2% agarose pad. Seam cells were visualized using the *scm::GFP* marker (*wIs51*) (Koh and Rothman, 2001) and the lateral side closer to the lens was counted on a Zeiss compound microscope (AxioScope A1) with 400x magnification.

RILs were phenotyped over a period of one week as they were growing at different rates. 116 RILs were scored twice and 38 RILs were scored thrice to make sure that RILs included in the pools for sequencing were the ones with the most reproducible seam cell number (SCN) between replicates. We pooled DNA of the two extreme groups (22 RILs in the low-SCN bulk and 24 RILs in the high-SCN bulk) for QTL mapping.

Single molecule mRNA fluorescent *in situ* hybridisation was performed using a Cy5-labelled *elt-6* and *hsp-110* probes (Biomers). Z-stacks with 17-30 slices, each of 0.7 μm, were acquired with a 100× oil immersion objective using an Andor iKon M 934 CCD camera system on a Nikon Ti Eclipse epifluorescence microscope. Region of interests for spot quantifications were drawn manually around seam cells visualized using the *scm::GFP* marker and a custom script in MATLAB was used to quantify mRNA molecules (Hintze et al., 2020).

Confocal images were obtained on a Leica SP5 using an Argon 488nm laser and analysed with Fiji. A region of interest was drawn from the vulva to the tail containing intestinal, vulval and seam cells expressing the POPHHOP marker. The corrected total cell fluorescence (CTCF) was calculated as follows: CTCF = Integrated Density - (Area of ROI × Mean fluorescence of background readings)

### Quantitative genetics

Recombinant inbred lines (RILs) were generated by crossing hermaphrodites from strain MBA256 (CB4856) and males from strain MBA231 (CB4856). F1 males from the cross were crossed to hermaphrodites from strain MBA290 (N2) containing the *egl-18(ga97)* mutation. Multiple new F1 hermaphrodites, which were egg-laying defective and carried the transgene *vtIs1[dat-1::gfp] wIs51[scm::GFP]* V were picked and allowed to self as cross progeny. 117 F2s which were egg-laying and carried only *wIs51[scm::GFP]* V transgene were picked onto single NGM plates and allowed to self. One hermaphrodite was transferred to a new plate for 10 – 14 generations to establish RILs. RIL-17 did not propagate during the selfing process so 116 RILs were produced in total.

We used a bulk segregant analysis approach to discover quantitative trait loci as previously done (Frezal et al., 2018). DNA was extracted from the low and high bulks using a Gentra Puregene Kit (Qiagen^®^). Whole genome sequencing was performed in an Illumina^®^ Hiseq platform (read length = 2 × 125), aiming to obtain at least 10 million read pairs per sample (25 × coverage). The whole genome sequencing data were analysed through the CloudMap Hawaiian variant mapping pipeline on a local galaxy server maintained in the lab (Minevich et al., 2012). The genotype for low-SCN and high-SCN was extracted into a single vcf file from the whole genome data using vcf combine on galaxy. The combined vcf was converted to tabular format using GATK tools on a bash terminal. Frequency of CB4856-like variation at each SNP position was calculated as the ratio of read counts with CB4856 SNP divided by the total number of reads. For the low-SCN and high-SCN bulk, the SNP positions where the genome quality (as determined by GQ scores < 40) for the sequencing was low were discarded using the QTLseqr package in R (Mansfeld and Grumet, 2018). SNPs where the total read depth was lower than 22 in the low-SCN bulk and lower than 24 in the high-SCN bulk were discarded as in (Frezal et al., 2018). We used log-odds ratio, which is the logarithm (base 10) likelihood ratio as a test statistic to evaluate if the deviations observed in SNP frequencies between the low-SCN and high-SCN bulk were statistically significant than expected from a null distribution. We calculated observed log-odds ratio using the following formula: 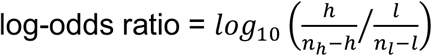, where l and h are mean SNP frequencies of CB4856 in a 300 kb window with no overlap multiplied by n_l_ and n_h_, n_l_ and n_h_ are number of RIL lines pooled for low-SCN and high-SCN bulks, respectively. The log-odds ratios under the null hypothesis were calculated where SNP frequencies for both the bulks were generated by 1 million simulated Bernoulli trails with p = q = 0.5. From these log-odds ratios, we found the two-tailed genome-wide threshold at a significance level (α = 0.05). The observed log-odds ratio was plotted against the genome location, and was compared to the genome-wide thresholds. The presence of a QTL was inferred when the log-odds ratio exceeded the threshold.

### CRISPR-Cas9 mediated genome editing

*hsp-110* and *sor-1* genetic variants in the N2 background were generated via injection of CAS9 ribonucleoprotein complexes (Paix et al., 2015) at the following final concentrations in 10 μl: tracRNA (0.75 nmol – IDT), custom crRNAs (*hsp-110* target sequence–AAA GTC AAT GGA GCA CGA CC and AAC CTT CTG ATT ACC ACC AT), (*sor-1* – TAG TCA ACG CCC ACC AAG CA, 20 μM, IDT) and Cas9 (1 μg/μl - IDT). The co-injection markers *myo-2::RFP* at 5 ng/μl and *rol-6(su1006)* at 40 ng/μl were used to select transgenic animals, and repair templates (6 μM) introducing SmaI sites in the case of *hsp-110* and ClaI in the case of *sor-1* together with the desired CB4856 SNP changes. F1 animals were screened for the co-injection markers, allowed to lay progeny and screened using restriction digests for the introduced sites. Positive lines were Sanger-sequenced to identify lines homozygous for SNP insertions. The co-CRISPR strategy (Arribere et al., 2014) was used to edit *bro-1*. An sgRNA targeting the following sequence (5’-AATCAATATACCTGTCAAGT) was cloned into pU6::unc-119 sgRNA vector by replacing the *unc-119* sgRNA as previously described (Friedland et al., 2013). The recovered icb45 allele is an in-frame deletion of 9 bp (GGAATCAATATA---------TTGGAATGGT) within the second exon of *bro-1*.

## Supplemental Figure Legends

**Figure S1:**
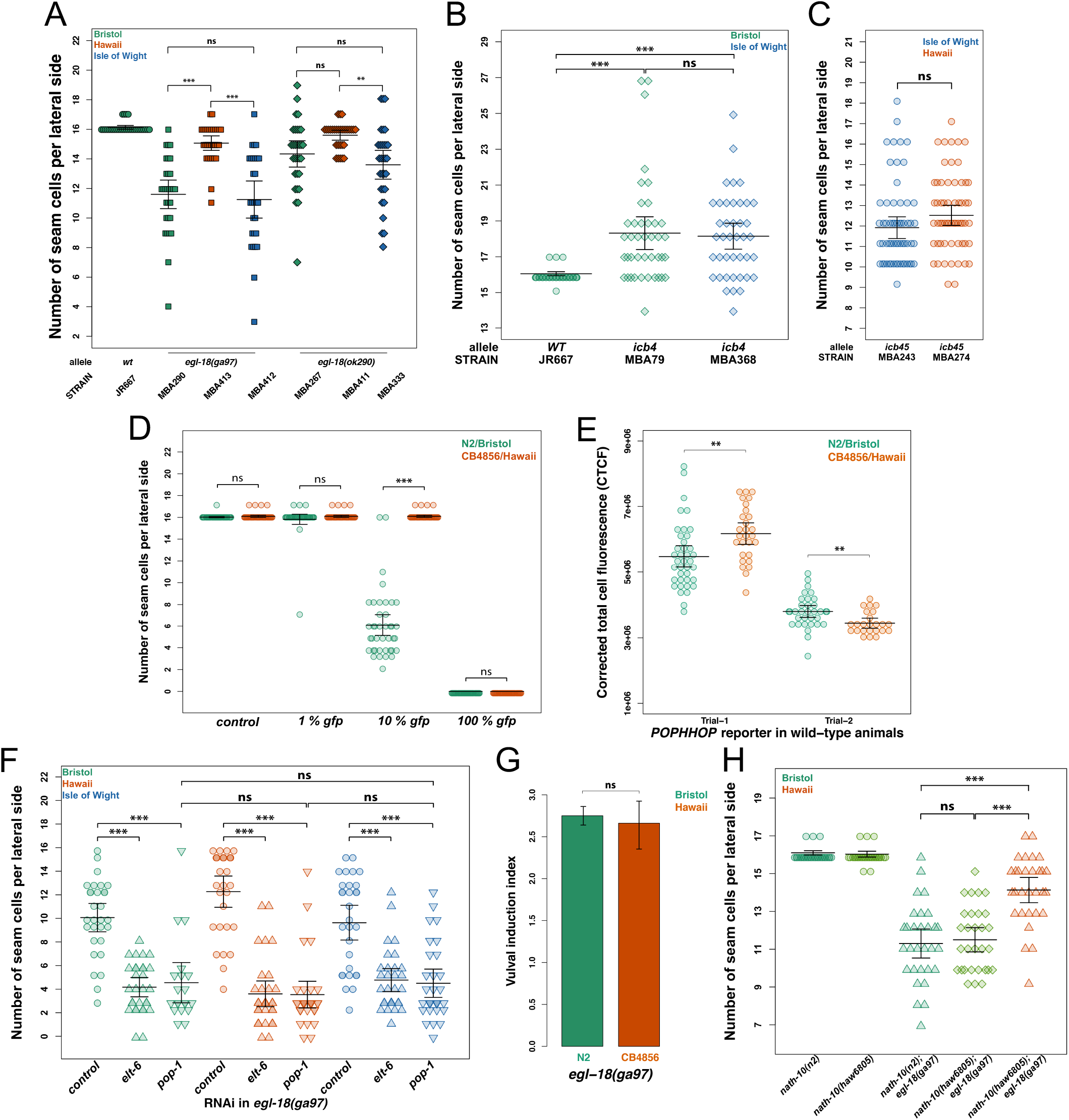
Differential mutation expressivity in the seam is specific to *egl-18* and requires strong *loss-of-egl-18* function and Wnt signaling. (A) The differential expressivity of *egl-18(ga97)* between N2 and CB4856 is less pronounced in the weaker allele *egl-18(ok290)*. There was no statistically significant difference in SCN between CB4856 and N2 (*p* = 0.06), but there was a significant difference in SCN between CB4856 and JU2007 carrying the *ok290* mutation (*p* = 0.0012), n ≥ 28 per strain. (B-C) No difference was found in the expressivity of *bro-1* or *eff-1* mutations between wild isolates. There is no statistically significant difference in SCN between the strains MBA79 (N2) and MBA368 (CB4856) carrying the same putative null allele *icb4* in *eff-1, n ≥* 37 per strain. Similarly, there is no statistically significant difference in SCN between the strains MBA274 (CB4856) and MBA243 (JU2007) carrying the same deletion allele *icb45* of *bro-1*, n = 60 per strain. (D) GFP RNAi is used to assess the efficacy of seam cell RNAi in CB4856 and N2 based on *scm::GFP* expression. A significant difference is revealed in 10% dilutions of the GFP dsRNA bacteria with control bacteria (HT115), but not in 1% dilution of GFP bacteria or when applied non-diluted (100% GFP), n = 40 per strain. (E) Quantification of corrected total fluorescence (CTCF) in N2 and CB4856 wild-type animals carrying the *POPHHOP* reporter. No consistent difference was found among different experimental trials, n ≥ 24 per strain (F) Knockdown of *elt-6* or *pop-1* by RNAi in *egl-18(ga97)* animals masks the differential expressivity observed between wild isolates. One-way ANOVA showed that SCN was not affected by strain upon *elt-6* RNAi (F (2, 87) = 1.52, *p* = 0.22), n = 30 per strain or *pop-1* RNAi (F (2, 77) = 0.86, *p* = 0.43), n ≥ 20 per strain (G) No significant difference was found in the vulval induction index between N2 and CB4856 carrying the *egl-18(ga97)* mutation, (n > 31 per strain, t-test). (H) The differential expressivity of the *egl-18(ga97)* mutation is not dependent on a previously known polymorphism in *nath-10*. There is a significant effect of strain on SCN (one-way ANOVA F (4, 145) = 77.19, *p* < 2.2 x 10^−16^). Post hoc Tukey HSD shows that there is a significant difference between *nath-10(haw6805); egl-18(ga97)* between N2 and CB4856 (*p* = 0). There was no difference between *nath-10(haw6805); egl-18(ga97)* and *nath-10(N2); egl-18(ga97)* (*p* = 0.98). n = 30 per strain. In all panels, error bars indicate 95 % confidence intervals and *** represent *p* value < 0.0001, ** *p* < 0.001 or * *p* < 0.05 by post hoc Tukey HSD test.

**Figure S2:**
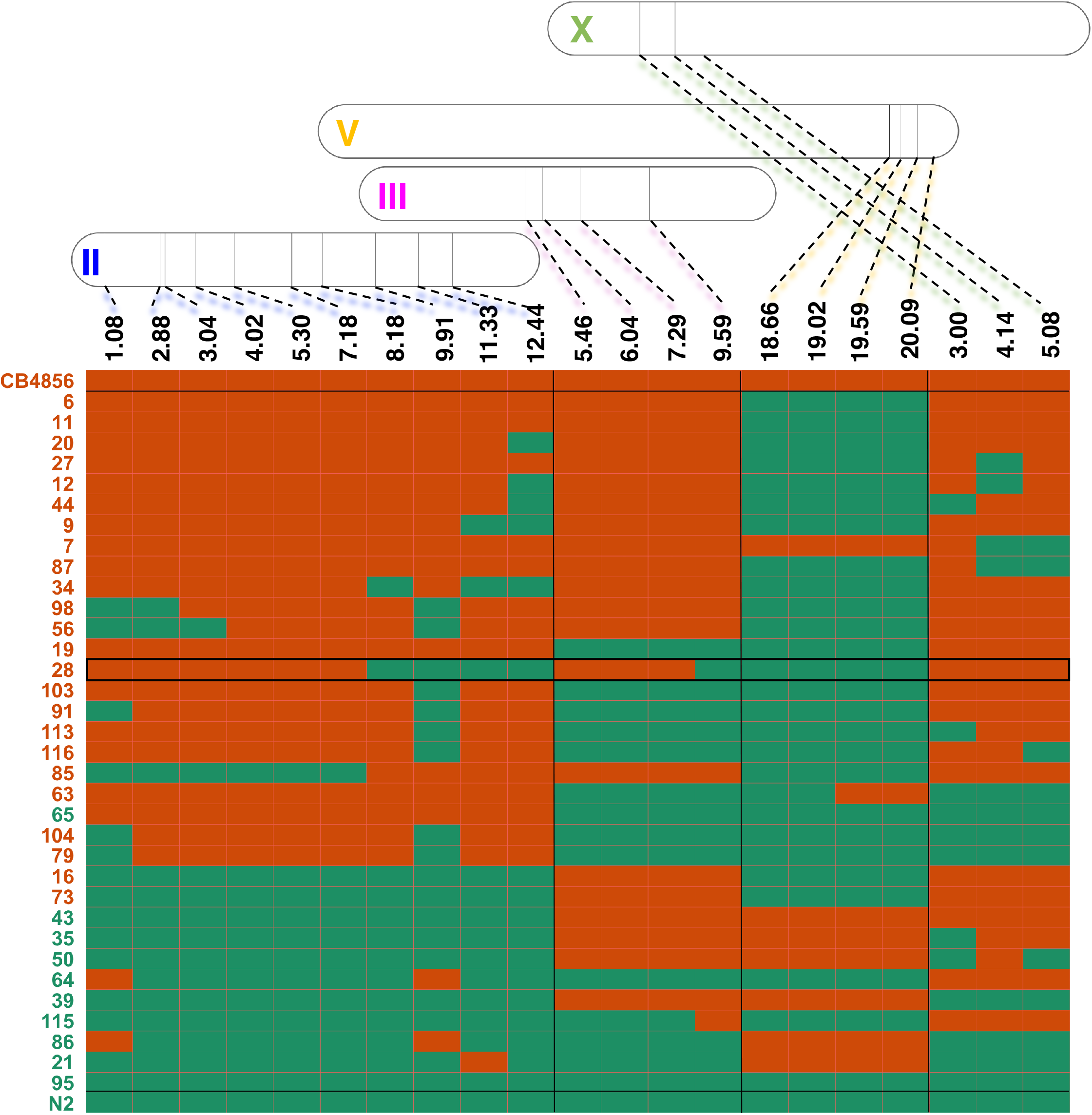
Genotyping of the individual RILs included in the low- and high-bulks using genetic markers. Each row represents an individual RIL line and its column a genetic marker on four different chromosomes to infer genetic composition per marker and per RIL. The location of the genetic markers on the chromosomes is shown above the graph. Markers on chromosome II, III, V and X are shown in blue, pink, yellow and green, respectively. Green and orange tiles represent N2 and CB4856 genomes respectively. RILs are ranked in descending order based on number of CB4856 genomic fragments they carry. RIL numbers are colour-coded based on whether they were included in the low-bulk (N2-like, green) or high-bulk (CB4856-like, orange). RIL-28, which was used for producing near isogenic lines containing QTLs on chromosomes II, III and X is highlighted in the middle.

**Figure S3:**
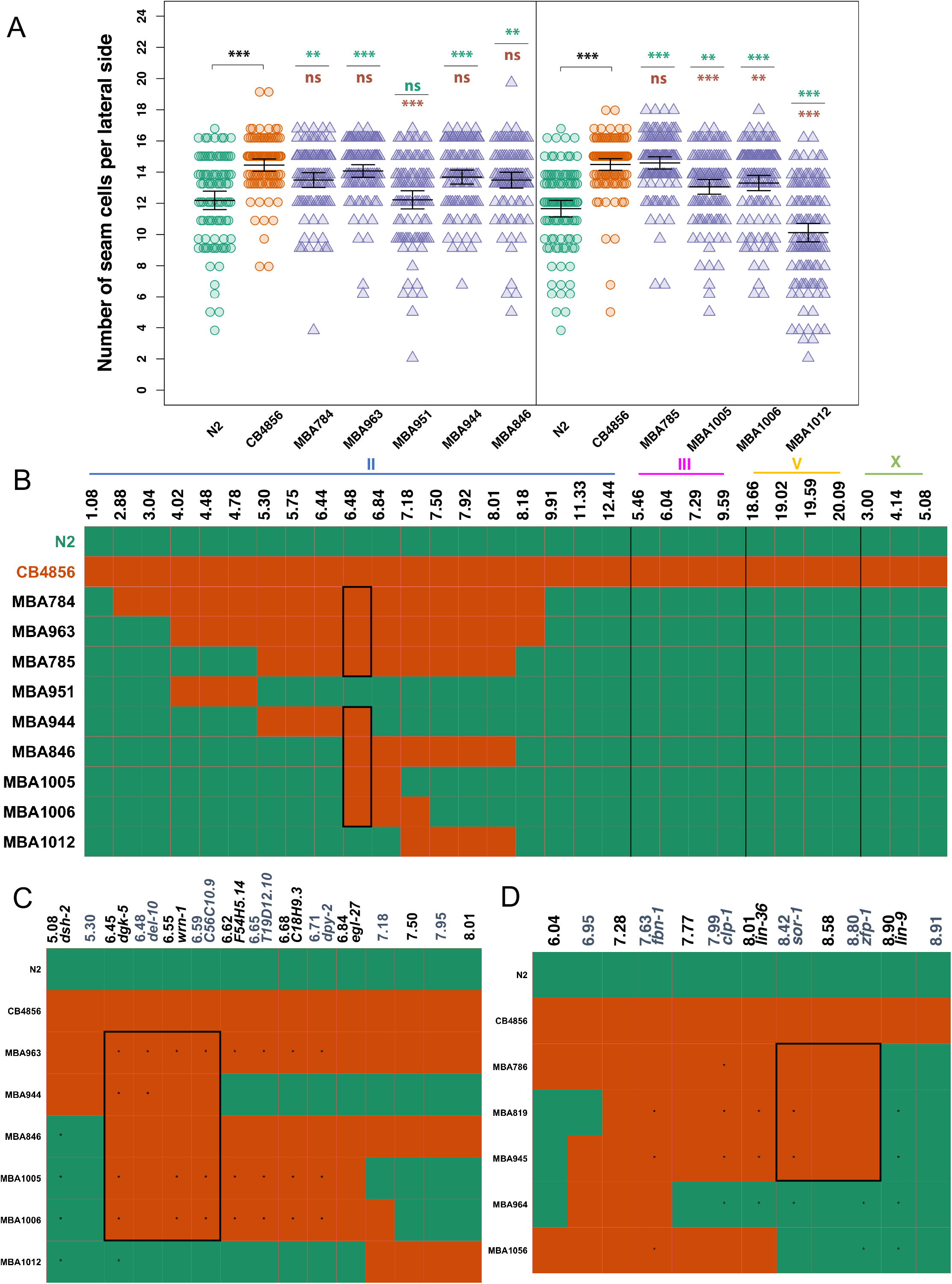
Genotype-to-phenotype analysis of NILs carrying genomic fragments of chromosome II from CB4856. (A) Seam cell number in near isogenic lines containing various fragments of chromosome II from CB4856 in the *egl-18(ga97)* N2 background. Vertical black line in the graph distinguishes between two independent experiments. In both experiments, one-way ANOVA showed that SCN was significantly affected by the strain (F (6, 635) = 12.25, p < 4.5 x 10^−13^; F (5, 622) = 51.18, *p* < 2.2 x 10^−16^, n ≥ 90 animals per strain. Error bars indicate average 95 % confidence intervals and *** *p* < 0.0001 or ** *p* < 0.001 correspond to significant differences by post hoc Tukey HSD compared to N2 (green stars) and CB4856 (orange stars) respectively. (B) Genotyping of NILs carrying various genomic fragments of chromosome II from CB4856. Each row represents a RIL line and its column its genetic composition at specific genetic markers, the location of which is shown above the graph. Green and orange tiles represent N2 and CB4856 genomes, respectively. Black-box in the middle highlights the genomic region that is common to the NILs that convert N2-like to CB4856-like *egl-18(ga97)* phenotype and absent in the NILs that do not. (C) Close-up on chromosome II showing the region that is common between the NILs that convert N2 *egl-18(ga97)* mutant expressivity to CB4856 levels (left). A close up of the region on Chr III, which includes *sor-1* and *hsp-110* is shown for comparison (right). In both cases, stars indicate inferred genotype for that marker.

**Figure S4:**
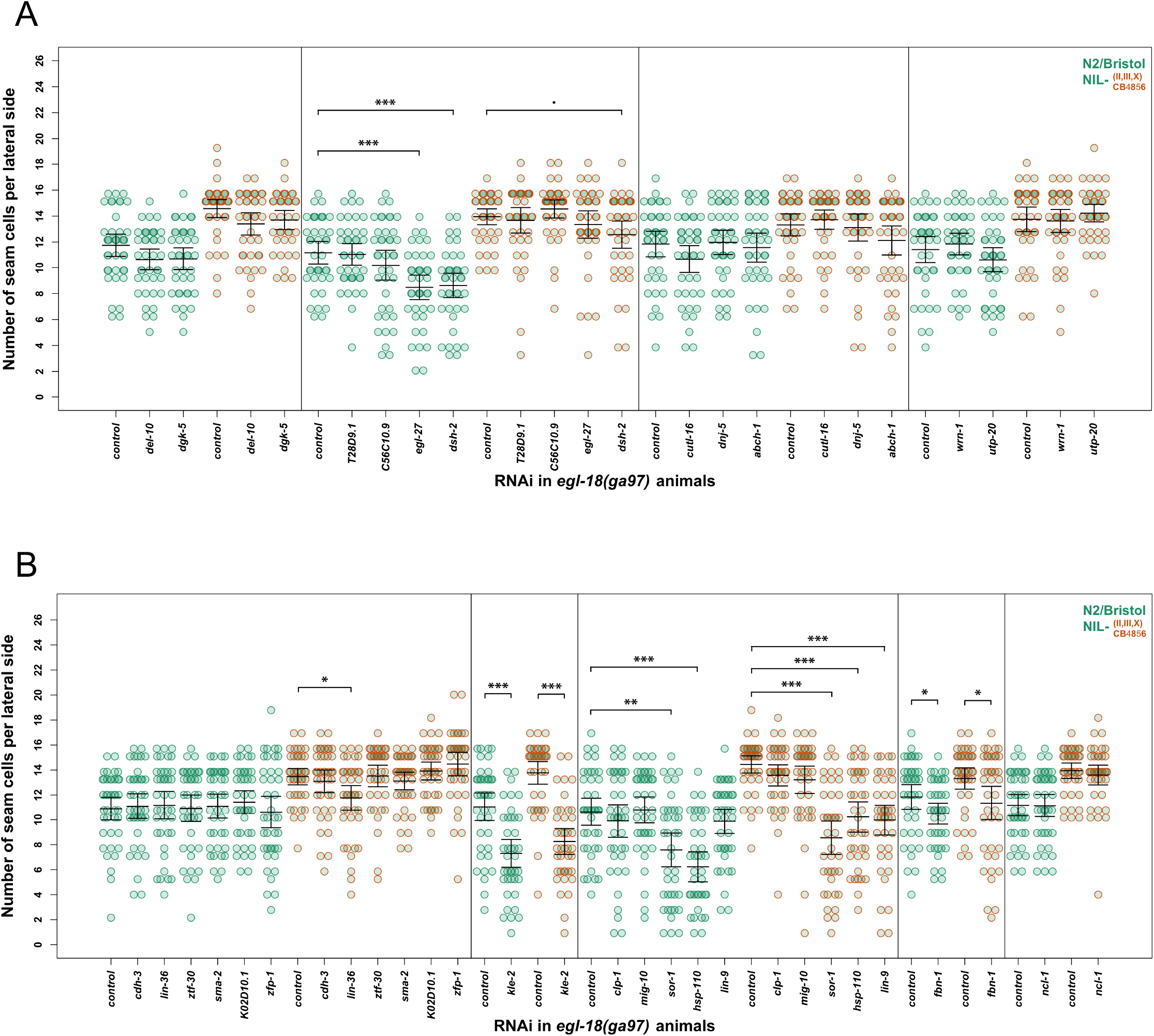
RNAi screen to identify candidate genes in the extended QTL regions of chromosomes II and III. (A-B) Seam cell number in mutant N2 and MBA848 (NIL carrying the QTLs on chromosome II, III and X) upon RNAi knockdown of candidate genes. (A) Knockdown of most genes on chromosome II did not have a significant effect on the SCN in mutant in N2 and MBA848, except for two genes (*egl-27* and *dsh-2*), which both had a significant effect. (B) Knockdown of two genes (*sor-1* and *hsp-110*) had a significant effect on the SCN in N2 and MBA848 (F (5, 234) = 10.48, p = 8.37 x 10^−9^; F (5, 234) = 19.65, p = 2.47 x 10^−16^, respectively). SCN decreased upon knockdown of *sor-1* and *hsp-110* in both N2 and MBA848 (*p* < 0.001). Knockdown of *lin-9* and *lin-36* showed a significant decrease in SCN only in MBA848. There was a decrease in SCN upon knockdown of *fbn-1* and *kle-2* in both N2 and MBA848. However, these genes were outside the final QTL interval. Error bars indicate standard deviation, n = 40 per strain and *** *p* < 0.0001, ** *p* < 0.001, * *p* < 0.05 correspond to significant differences compared to control RNAi by ANOVA and post hoc Dunnett’s multiple comparison test. In both A and B, dashed lines mark independent experiments therefore control treatments are repeated.

## Supplementary Tables

**Table S1. Strains used in this study.**

**Table S2. Primers used in this study (general oligos, genotyping and smFISH).**

**Table S3. Genetic variants within QTL regions on chromosome II and III**

## Acknowledgments

We thank Mingke Pan for technical help with genotyping. Some *C. elegans* strains were provided by the CGC, which is funded by NIH Office of Research Infrastructure Programs (P40 OD010440). This work was funded by the by the European Research Council (ROBUSTNET).

